# *CDKN2C* homozygous loss identifies a distinct subtype of *TP53/RB1*-wildtype leiomyosarcoma with frequent *CIC* genomic alterations and 1p/19q-codeletion

**DOI:** 10.1101/2020.03.02.973305

**Authors:** Erik A. Williams, Radwa Sharaf, Brennan Decker, Adrienne J. Werth, Helen Toma, Meagan Montesion, Ethan S. Sokol, Dean C. Pavlick, Nikunj Shah, Jeff M. Venstrom, Brian M. Alexander, Jeffrey S. Ross, Lee A. Albacker, Douglas I Lin, Shakti H. Ramkissoon, Julia A. Elvin

**Affiliations:** Foundation Medicine, Inc., 150 Second Street, Cambridge, Massachusetts 02141; Department of Pathology, Brigham and Women’s Hospital and Harvard Medical School, Boston, Massachusetts; Christiana Hospital, Department of Obstetrics and Gynecology, 4755 Ogletown Stanton Rd, Newark, DE 19713; Department of Pathology, State University of New York Upstate Medical University, 766 Irving Avenue, Syracuse, New York 13210; Wake Forest Comprehensive Cancer Center and Department of Pathology, Wake Forest School of Medicine, Winston-Salem, North Carolina, 27157

**Keywords:** Uterine leiomyosarcoma, comprehensive genomic profiling, *CDKN2C*, *CIC*, 1p/19q-codeletion

## Abstract

**Purpose:** Leiomyosarcomas (LMS) harbor frequent inactivation of *TP53* and *RB1*, and extensive DNA copy number alterations. Here, we describe a distinct recurrent genomic signature in *TP53*/*RB1*-wildtype uterine LMS.

**Methods:** Tissues from 276,645 unique advanced cancers, including 2,570 uterine and soft tissue LMS were sequenced by hybrid-capture-based next-generation DNA and RNA sequencing/comprehensive genomic profiling of up to 406 genes. We characterized clinicopathologic features of relevant cases.

**Results:** We identified 77 LMS with homozygous copy loss of *CDKN2C* at chromosome 1p32.3 (3.0% of LMS). Genomic alterations (GAs) in *TP53, RB1*, and *ATRX* were rare in comparison with the remainder of the LMS cohort (11.7% vs 73.4%, 0% vs 54.5%, 2.6% vs 24.5%, all p<0.0001). *CDKN2C*-null LMS cases were significantly enriched for GAs in *CIC* (40.3% vs 1.4%) at 19q13.2, *CDKN2A* (46.8% vs 7.0%), and *RAD51B* (16.9% vs 1.7%; all p<0.0001). Chromosome arm-level aneuploidy analysis of available LMS cases (n=1,251) found that 85% (n=33/39) of *CDKN2C*-null LMS cases exhibited 1p/19q-codeletion, with significant enrichment in comparison to the remainder of the evaluated LMS cohort (85% vs. 5.1%, p<0.0001).

99% of *CDKN2C*-null cases were in females; median age was 61 years at surgery (range, 36-81 years). 55 cases were of uterine primary, 4 cases non-uterine, and the remaining 18 of uncertain primary site. 6 patients had a prior history of leiomyomatosis or uterine smooth muscle tumor of uncertain malignant potential. 60% of cases showed at least focal epithelioid variant histology. Most cases were of known advanced stage, with 62% of confirmed uterine primary cases at FIGO stage IVB.

**Conclusion:** The identification of this novel genomic subset may have prognostic and/or therapeutic clinical relevance, including use of specific cyclin-dependent kinase inhibitors.

## Main

Leiomyosarcoma (LMS), a neoplasm defined by smooth muscle differentiation, is the most common form of uterine sarcoma.^1^ LMS is aggressive and challenging to treat given its resistance to standard therapy, as documented by high rates of recurrence and progression. Multiple studies have shown an overall 5 year survival of 25 to 76%, with survival for women with metastatic disease at presentation approaching 10-15%.^2^ Stage of disease, as defined by the International Federation of Gynecology and Obstetrics (FIGO)^3^ or the American Joint Committee on Cancer (AJCC)^4^, at the time of diagnosis is the most important prognostic factor for uterine LMS.^1^ Surgery is the standard of care for localized tumors, with hormonal and cytotoxic chemotherapy reserved for advanced stages.^5^

Genomic studies of LMS have demonstrated significant mutational heterogeneity, frequent inactivation of *TP53* and *RB1* through varied mechanisms, and widespread copy number alterations.^6^ LMS is often associated with complex karyotypes with numerous chromosomal gains and losses.^7^ LMS has demonstrated occasional potentially targetable genomic alterations (GAs), but novel targeted therapeutic agents have not been widely used.^8,9,10^ Herein, we describe a novel recurrent genomic signature of *CDKN2C* homozygous loss in leiomyosarcoma primarily from the uterus, with significantly low frequency of *TP53* and *RB1* genomic alterations.

## Methods

### Cohort

Comprehensive genomic profiling (CGP) was performed in a Clinical Laboratory Improvement Amendments (CLIA)-certified, CAP (College of American Pathologists)-accredited laboratory (Foundation Medicine Inc., Cambridge, MA, USA). Approval for this study, including a waiver of informed consent and a HIPAA waiver of authorization, was obtained from the Western Institutional Review Board (Protocol No. 20152817). The pathologic diagnosis of each case was confirmed on routine hematoxylin and eosin (H&E)-stained slides. In brief, ≥60 ng DNAs was extracted from 40 μm of 276,645 cancer specimens, including 2,570 leiomyosarcoma specimens and 19 leiomyomatosis cases, in formalin-fixed, paraffin-embedded tissue blocks. The samples were assayed by CGP using adaptor ligation, and hybrid capture was performed for all coding exons from 287 (version 1) to 315 (version 2) cancer-related genes plus select introns from 19 (version 1) to 28 (version 2) genes frequently rearranged in cancer (Supplemental Table 1). Various samples were similarly assayed but performed in DNA on 406 genes and selected introns of 31 genes involved in rearrangements (Supplemental Table 1), and in RNA on 265 genes commonly rearranged in cancer. Sequencing of captured libraries was performed using the Illumina HiSeq technology to a mean exon coverage depth of > 500×, and sequences were analyzed for all classes of genomic alterations including short variant alterations (base substitutions, insertions, and deletions), copy number alterations (focal amplifications and homozygous deletions), and select gene fusions or rearrangements.^11–13^ To maximize mutation detection accuracy (sensitivity and specificity) in impure clinical specimens, the test was previously optimized and validated to detect base substitutions at a ≥5% mutant allele frequency (MAF), indels with a ≥10% MAF with ≥99% accuracy, and fusions occurring within baited introns/exons with > 99% sensitivity.^11^ Tumor mutational burden (TMB, mutations/Mb) was determined on 0.8-1.1 Mbp of sequenced DNA.^13^ Microsatellite instability (MSI) was determined on up to 114 loci.^14^

**Table 1.** Clinical characteristics of patients with leiomyosarcoma with homozygous loss of *CDKN2C.*

| Characteristic | No. (%) |
| --- | --- |
| No of patients | 77 |
| Median age at diagnosis, years (range) | 61 (36-81) |
| Sex |  |
| Female | 76 (99) |
| Male | 1 (1) |
| Primary site |  |
| Uterine | 55 (71) |
| Soft tissue | 4 (5) |
| Indeterminant | 18 (23) |
| FIGO staging (uterine primary) |  |
| IB | 6 (11) |
| IIA | 2 (4) |
| IIB | 4 (7) |
| IIIA | 2 (4) |
| IIIB | 1 (2) |
| IIIC | 4 (7) |
| IVB | **34 (62)** |
| Unknown | 3 (6) |
| AJCC staging (soft tissue or indeterminant primary) |  |
| IA | 1 (5) |
| IV | **19 (86)** |
| Unknown | 2 (9) |

### Copy Number Analysis

Copy number analysis to detect gene-level amplifications and homozygous deletions was performed as previously described.^11^ In short, each specimen was analyzed alongside a process-matched normal control, which custom algorithms used to normalize the sequence coverage distribution across baited DNA regions. Normalized coverage data and minor allele SNP frequencies were fit by statistical models which minimized error in purity and ploidy dimensions.

Signal-to-noise ratios for each genomic segment were used to determine gain or loss status; the sum of those segment sizes determined the fraction of each arm gained or lost. Chromosomes were assessed for arm level aneuploidy, defined as positive if >50% of the arm was altered. This threshold was previously validated on 109 *IDH1*/*2*-mutant glioma samples with 1p/19q-codeletion FISH results available. Cases were blinded to FISH results and 1p/19q-codeletion status was determined via arm level aneuploidy analysis. Concordance was 95%, sensitivity 91%, and positive predictive value was 100%. A query for chromosome 1p and 19q arm level aneuploidy was performed on LMS cases with available aneuploidy data (n=1,251), with positive cases defined as 1p/19q-codeleted.

### Clinicopathological Analysis of Leiomyosarcoma Cohort Harboring CDKN2C Inactivation

The cohort of leiomyosarcomas comprised 77 cases, each from a different patient, that were submitted for CGP (Foundation Medicine Inc., Cambridge, MA, USA) during routine clinical care. Human investigations were performed after approval by a local human investigations committee and in accordance with an assurance filed with and approved by the Department of Health and Human Services, where appropriate. Clinicopathological data including patient age, gender, tumor site, and FIGO stage or AJCC stage (8^th^ edition) were extracted from the accompanying pathology report.^4,15^ Primary site data was not available for a subset of cases (“indeterminant primary”). The histopathology was assessed on routine H&E-stained slides of tissue sections submitted for genomic profiling by two board-certified pathologists (E.A.W., D.L.).

Quantitative data were analyzed using the Fisher exact test owing to the categorical quality of the data and the size of the cohort. For age and TMB comparison between two groups, the non-parametric Mann Whitney U test was used. A two□tailed *P*□value of <0.05 was considered to be statistically significant.

## Results

### Clinicopathologic Features

From an internal series of 276,645 unique advanced cancers, including 2,570 leiomyosarcomas (LMS), of which 939 were of confirmed uterine origin, we identified 77 confirmed LMS with homozygous copy loss of *CDKN2C* at chr 1p32.3 (3.0% of all LMS, 5.9% of uterine LMS). Clinical characteristics are summarized in Table 1. Patients were significantly older than the remainder of the LMS cohort (median age 61 vs. 57 years, p=0.0009, Mann Whitney U test). Patients were enriched for female gender in comparison to the remainder of the LMS cohort (99% vs. 79% [1968/2493], p<0.0001, Fisher’s test). 6 female patients had a prior history of leiomyomatosis (n=2) or uterine smooth muscle tumor of uncertain malignant potential (n=4). The majority of patients showed clinically advanced/metastatic disease, with 62% of confirmed uterine primary cases documented at FIGO stage IV, and 86% of indeterminant or soft tissue primary cases at AJCC stage IV (Table 1).

29 cases were sequenced from tumor at primary site [25 uterine, one primary abdominal wall, one primary hip/gluteal, one primary sacrum, and one primary small bowel]. The remaining 48 cases were sequenced from tumor to metastatic sites, including lung (n=7), retroperitoneum (n=5), abdominal wall (n=4), limb soft tissue (n=4), omentum (n=4), liver (n=3), paraspinal (n=3), peritoneum (n=3), pleura (n=2), colon (n=2), kidney (n=2), chest wall (n=2), mesentery (n=2), small intestine (n=1), hilar lymph node (n=1), vagina (n=1), posterior mediastinum (n=1), and heart (n=1).

### Comprehensive genomic profiling

The distribution of genomic alterations (GA) of the 77 cases is displayed in Figure 1. The rate of oncogenic mutations for genes of interest was compared between *CDKN2C*-null cases and the remainder of the LMS cases, with p values (Table 2).

**Table 2.** Percent frequency of genomic alterations by *CDKN2C* status, with p values.

| Genomic alteration | *CDKN2C*- null LMS (%) | Remaining LMS cohort (%) | *P* |
| --- | --- | --- | --- |
| *TP53* | 11.7 | 73.4 | <0.0001 |
| *RB1* | 0 | 54.5 | <0.0001 |
| *ATRX* | 2.6 | 24.5 | <0.0001 |
| *PTEN* | 9.1 | 16.0 | p=0.113 |
| 1p/19q-codeletion | 84.6 | 5.1 | <0.0001 |
| *CIC* | 40.3 | 1.4 | <0.0001 |
| *CDKN2A* | 46.8 | 7.0 | <0.0001 |
| *RAD51B* | 16.9 | 1.7 | <0.0001 |

Abbreviations: LMS, leiomyosarcoma

**Figure 1.**
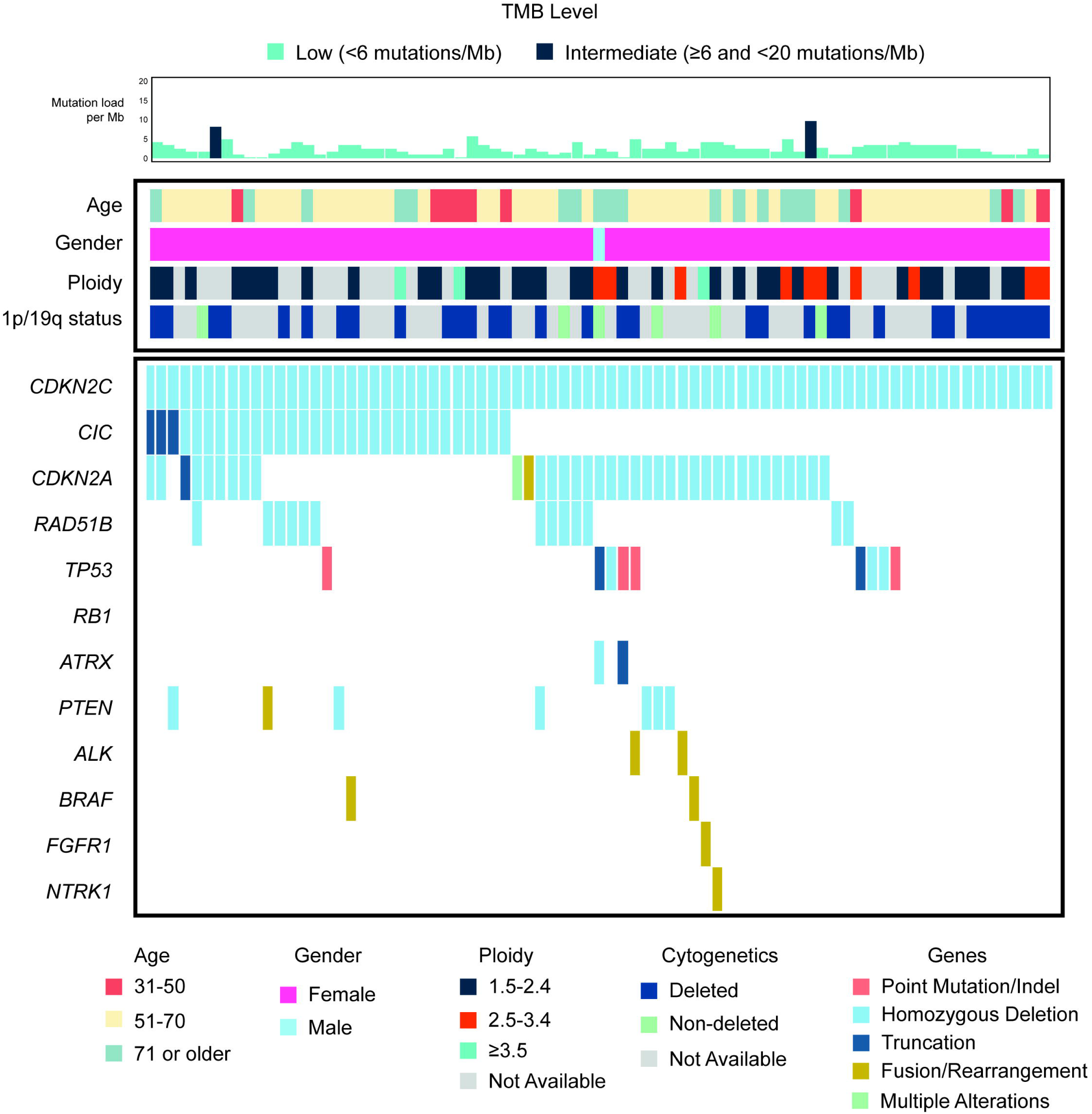
Molecular profiles of *CDKN2C*-null leiomyosarcoma.

Among the 77 *CDKN2C*-null LMS, *TP53, RB1*, and *ATRX* genomic alterations (GAs) were rare in comparison to the remainder of the LMS cohort. *CDKN2C*-inactivated LMS comprised 14% of *TP53*/*RB1*-wildtype LMS (68 of 486). The cases were also significantly enriched for GAs in *CIC* at 19q13.2, as well as *CDKN2A* and *RAD51B* (Table 2). GAs in *CIC* and *CDKN2A* were predominantly deep deletions and exclusively deep deletions for *RAD51B*. Additional variants included truncating variants in *CIC* (n=3 cases) and inactivating structural rearrangements in *CDKN2A* (n=2 cases). Rare activating fusion events were also identified in *ALK* (2 cases), *BRAF* (2 cases), *FGFR1* (1 case), and *NTRK1* (1 case) (Fig 1). 85% of cases showed deep deletion of *FAF1* at 1p32.3, a gene adjacent to *CDKN2C* [9.7 kb apart].

Median TMB was 2.4 mut/Mb (range <0.8-9.6; Q1-Q3=1.6-3.2), similar to the remainder of the LMS cohort (median=2.4 mut/Mb; range <0.8-203; Q1-Q3=1.6-4.0), but slightly lower overall (p=0.0425, Mann Whitney U test). No microsatellite unstable cases were present in the *CDKN2C*-null cohort.

3 patients had 2 separate tissue specimens analyzed (Supplemental Table 2). One patient had sequencing of both the primary uterine LMS, and a subsequent lung metastasis. The lung mass showed additional homozygous loss of *CIC*.

A query and review for chromosome 1p and 19q arm level aneuploidy in available LMS cases (n=1,251) revealed 100% of *CDKN2C*-null LMS cases to have whole-arm aneuploidy of the short arm of chromosome 1, and 85% to have aneuploidy of the long arm of chromosome 19 (1p/19q-codeletion) (n=33/39). Significant enrichment for 1p/19q co-deletion was identified in comparison to the remainder of the evaluated LMS cohort (85% vs. 5.1%, p<0.0001). Sample copy number plots of *CDKN2C*-null LMS exhibiting 1p/19q-codeletion are shown in Figure 2A,B. Additional recurrent chromosomal arm level changes were identified in the 39 cases available, including most frequently aneuploidy of chromosome 6p (n=18 cases), 9p (n=12), 10q (n=16), 11p (n=23), 13q (n=26), 14q (n=24), and 16q (n=26).

**Figure 2.**
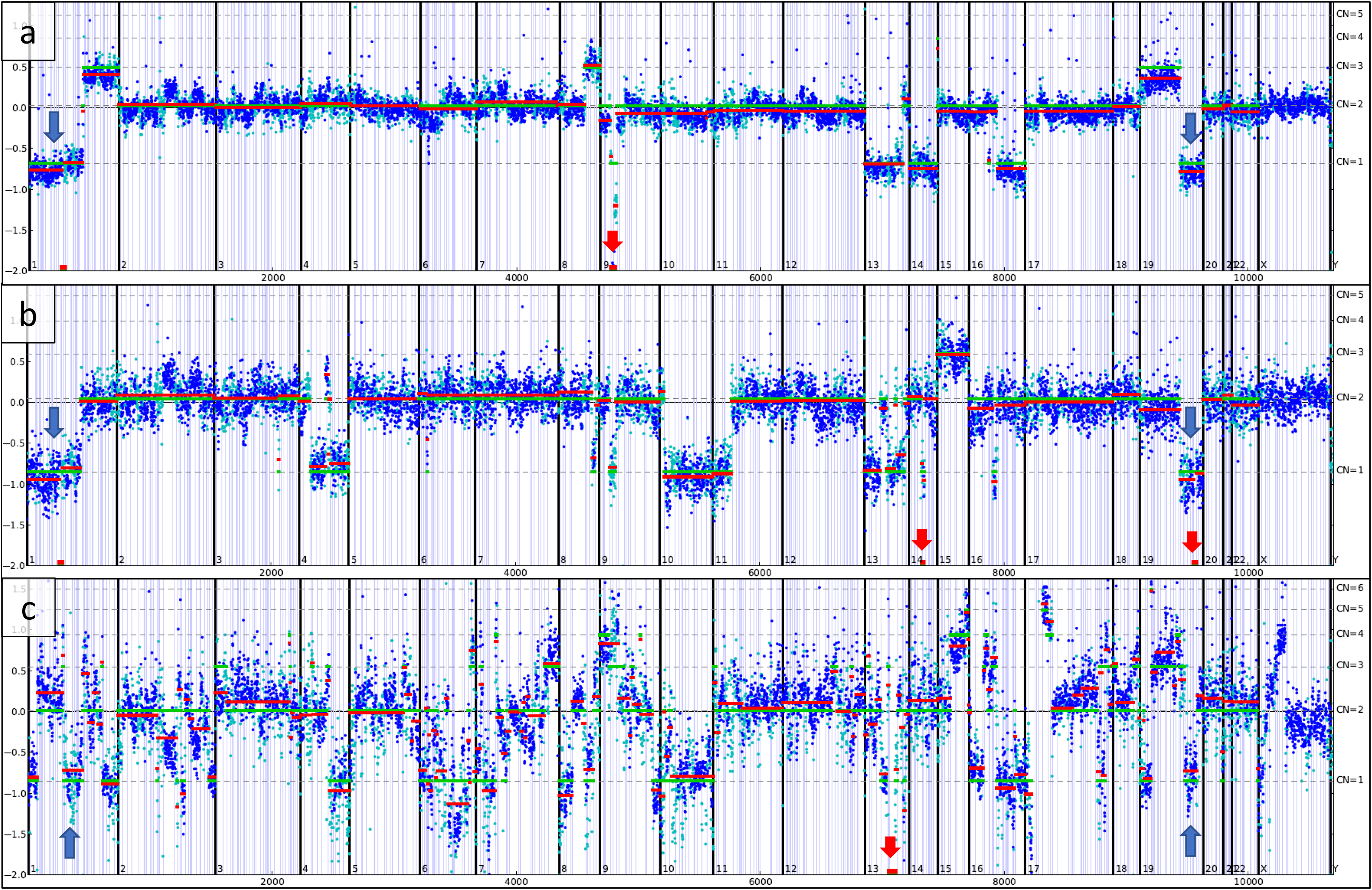
Copy number plots of leiomyosarcoma with 1p and 19q arm level aneuploidy. The Y axes display log-ratio measurements of coverage from each case as compared to a normal reference sample, with assessed copy numbers denoted by dashed horizontal lines. Each dot represents a genomic region evaluated by the assay (cyan=SNP, blue=exon), which are organized by genomic position. Red lines designate average log-ratio in a segment, while green lines represent model prediction. **(a)** *CDKN2C*-null leiomyosarcoma with a truncating variant in *CIC* (p.Q907*) and deep deletion of *CDKN2A* at chromosome 9p21.3. **(b)** *CDKN2C*-null leiomyosarcoma with homozygous deletion of *CIC* at chr. 19q13.2 and *RAD51B* at chr. 14q24.1. **(c)** *CDKN2C*-retained leiomyosarcoma with deep deletion of *RB1* at chr. 13q14.2 and a copy number plot of high complexity.

A review of 1p/19q-codeletion status in available LMS cases without *CDKN2C* deep deletion (n=1,212) revealed 62 1p/19q-codeleted cases (5.1%) (Fig 2C). The cases showed diverse GAs, including significant percentages of GAs in *TP53* (51.6%), *RB1* (45.2%), *ATRX* (16.1%), and *PTEN* (12.9%). GAs were also identified in *CDKN2A* (22.6%) and *ALK* (9.7%; all activating rearrangement events). A minority of cases showed GAs in *RAD51B* (8.1%), *CIC* (6.5%; all homozygous loss), and *FAF1* (5.2%). 3 of the 4 cases with deep deletion of *CIC* also showed deep deletion of both *RAD51B* and *FAF1*. All 4 occurred in uterine LMS [one with history of STUMP].

All non-LMS sarcoma cases (n=12,097) were also evaluated for *CDKN2C* homozygous loss. 22 of 1297 gastrointestinal stromal tumors (GISTs) had deep deletion of *CDKN2C* (1.7% of GISTs). 21 had a *KIT* mutation and the remaining case had a *PDGFRA* mutation; none of the 14 cases with 1p/19q data had 1p/19q-codeletion. 19 additional sarcoma cases with deep deletion of *CDKN2C* were identified (0.18% of non-LMS non-GIST sarcomas). These included diverse sarcoma diagnoses, including six high grade sarcomas NOS, two osteosarcomas, two malignant peripheral nerve sheath tumors, and two inflammatory myofibroblastic tumors. Genomics were also varied, with alterations identified in *CDKN2A* (68.4%), *TP53* (42.1%), *NF1* (26.3%), *NF2* (26.3%), and *ALK* (15.8%; all activating rearrangement events). No genomic alterations in *CIC* or *RAD51B* were identified. 11 of the 19 cases had 1p/19q-codeletion data available; 2 of the 11 (18%) had 1p/19q-codeletion. Both were *ALK*-rearrangement positive cases in females [ages 63 and 72 years]. Outside pathology report review revealed diagnoses of a uterine inflammatory myofibroblastic tumor and an ER-positive (by immunohistochemistry) recurrent unclassified intermediate-grade myxoid spindle cell sarcoma involving colon [primary site not available].

The LMS cohort was also evaluated for cases with pathogenic alterations in *CDKN2C* other than deep deletion. Only a single case was identified, in a 52-year-old female with an ER-positive, PR-positive (per report, by immunohistochemistry) uterine LMS with truncating mutation in *CDKN2C* (p.R68*). Concurrent deep deletion of *CIC* and *RAD51B* were identified; 1p/19q status was not available for the case.

### Histopathology

Histopathologic evaluation was performed on high-resolution digital pathology H&E slides of 70 cases. Histology was heterogeneous, as shown in Figure 3. 27 cases (39%) were epithelioid LMS. Of these cases, 2 showed follicle-like spaces, and a single case showed focal pleomorphic giant cells. 23 cases (33%) were spindle cell LMS. 19 cases (27%) showed mixed histology, including 11 mixed spindle and epithelioid LMS [one with focal pleomorphic giant cells], 4 mixed spindle and myxoid LMS, 2 mixed epithelioid and myxoid LMS, and 2 mixed spindle, epithelioid and myxoid LMS. A single case showed small round cell morphology.

**Figure 3.**
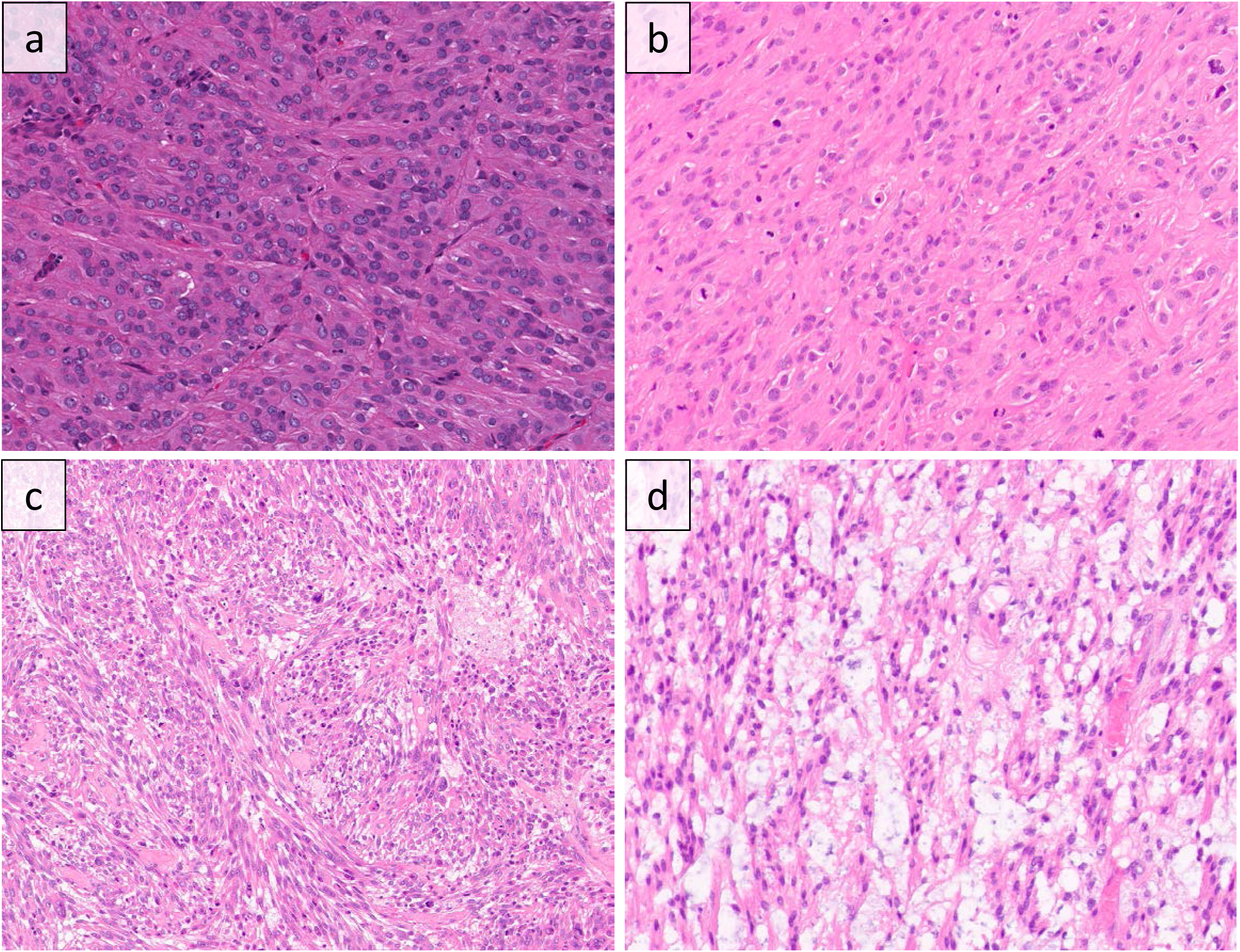
Histopathology of *CDKN2C*-null leiomyosarcoma ranged from epithelioid (**a**,**b**), to spindled (**c**) (H&E stains, 200x). Occasional cases showed focal myxoid histology (**d**) (H&E stain, 200x).

## Discussion

In the 2,570 cases of LMS evaluated, we discovered that cases with *CDKN2C* homozygous deletion (n=77; 3.0%) comprise a genomically distinct molecular subgroup. Cases lack mutations in *TP53, RB1*, and *ATRX*, but showed frequent 1p/19q-codeletion (85%), and close to half showed inactivation of *CIC*. The vast majority occur in females, and most are of uterine primary site of origin. A high percentage demonstrated epithelioid variant features on histology, and limited clinical data suggests a possible association with and progression from lower grade uterine smooth tumors, such as leiomyomatosis and smooth muscle tumor of uncertain malignant potential.

*CDKN2C* at 1p32.3 encodes the homologous p18INK4C cell cycle regulatory protein that blocks cell cycle progression by inhibiting cyclin D-dependent kinases CDK6 and CDK4.^16,17,18^ Loss of *CDKN2C* results in loss of potent inhibition of CDK4/6 in the cyclin D-CDK4/6-INK4-Rb pathway. CDKN2C is also a key factor for ATM/ATR-mediated p53 activation, and *CDKN2C* loss has been shown to block p53 induction in response to DNA damage.^19,20^ *CDKN2C* loss has been documented in a subset of diverse tumor types, including multiple myeloma, pituitary adenoma, and thyroid carcinoma.^21–23^ *CDKN2C* loss has also been documented in a small percentage of oligodendroglioma.^24,25^The adjacent *FAF1* gene at 1p32.3 encodes FAS associated factor 1, which enhances Fas-mediated apoptosis, and may contribute to tumor pathogenesis.^26^

The *CIC* gene on chromosome 19q13.2 is the human homologue of the *capicua* gene in Drosophila. In Drosophila, *capicua* encodes CIC, which represses genes induced downstream to RTK pathway activation.^27^ In the absence of RTK signaling, CIC blocks transcription of genes that have diverse effects on cellular proliferation, metabolism, and migration.^28^ Along with single copy loss of *CIC* on 19q, concurrent inactivating mutations in *CIC* are identified in a high percentage of oligodendrogliomas.^29^

1p/19q whole arm co-deletion, with concurrent mutation in *IDH1* or *IDH2*, are entity-defining alterations for oligodendroglioma.^30,31,32^ Oligodendrogliomas are typically associated with relatively long overall survival, and treatment strategies used are often stratified based on 1p/19q status.^33,34,35^ The co-deletion is a result of unbalanced translocation between two chromosomes with subsequent loss of der(1;19)(p10;q10), likely because chromosomes 1 and 19 are near each other in the non-random organization of the nucleus.^36,37,38^ A significant percentage of oligodendrogliomas also show *CIC* and *FUBP1* mutations.^28^ Our cohort of *CDKN2C*-null LMS shows notable similarities and differences to oligodendroglioma. 40% showed an inactivating alteration in *CIC*, most commonly homozygous deletion. While *FUBP1* at 1p31.1 is somatically mutant in a subset of oligodendroglioma, no *CDKN2C*-null LMS cases in our cohort had inactivating genomic alterations in *FUBP1*, or pathogenic alterations in *IDH1*/2 or *TERTp*. Rare sarcoma-like tumors originating from oligodendroglioma have been reported, with documented *IDH1* mutation and 1p/19q co-deletion, and have been termed “oligosarcoma”.^39,40,41^

Among all sarcomas with *CDKN2C* loss, 1p/19q-codeletion appears to be nearly exclusive to LMS. However, in the overall LMS cohort, occasional cases with *TP53* and *RB1* alterations were positive by the 1p/19q-codeletion detection algorithm. We speculate that given the complexity of these genomically-unstable cases, occasional *CDKN2C*-retained LMS cases satisfy the criteria for detection (Figure 2C). As such, identification of *CDKN2C* deep deletion may be the most specific distinguishing feature.

Cytogenetic findings in LMS and leiomyoma have been previously reported. A greater frequency of loss of 1p has been documented in metastasized LMS.^42^ From a cytogenetics study of 800 uterine leiomyomata, nine diploid cases with 1p loss were identified, with association with other alterations, particularly loss of chromosome 19 and/or 22.^43^ Transcriptional profiling of two of the 1p deleted leiomyomas in this study showed alignment with malignant LMS in a hierarchical clustering analysis.^43^ In another study, 1p loss was identified in approximately one quarter of uterine cellular leiomyomata.^44^ Three reports on a total of eight pulmonary-based “benign metastasizing leiomyoma” reported 19q and 22q terminal deletion in each case.^45–47^ Rare uterine leiomyomas with genomic alterations in *RAD51B* have also been identified.^48^ The overlap in GAs between our cohort of LMS and a subset of leiomyoma in the literature suggests a possible connection between these entities. The limited clinical history in our cohort further suggests a link with lower grade smooth muscle neoplasms.

Gene expression profiles have identified different molecular subtypes associated with distinct clinical outcomes.^49^ Distinct molecular subtyping based on differential expression of a large number of genes has been proposed.^49^ Future studies are needed to identify the gene expression profile of this uncommon genomic subtype.

Evaluation of genomic alterations in epithelioid or myxoid uterine smooth muscle neoplasms are limited in the literature.^50–54^ While a high percentage of *CDKN2C*-null LMS in our study demonstrated epithelioid features on histology, histology was also varied, and conclusions are limited. Immunohistochemistry results extracted from pathology reports were typical for uterine LMS, with expression of characteristic smooth muscle markers (data not shown).^55^

Given the overall low response rate of LMS to standard therapy, the identification of this targetable alteration in *CDKN2C* that defines a genomically distinctive subset of LMS may be useful for decisions on therapy. Well-known CDK4/6 inhibitors have previously shown effectiveness in a single case of LMS with a *CDKN2A* alteration.^10^ Nearly half of the *CDKN2C*-null LMS harbored loss of *CDKN2A*; CDK4/6 inhibitors may be effective in targeting both alterations for these cases. A subset of *CDKN2C*-null and/or 1p/19q co-deleted LMS also harbored activating fusions in *ALK*, for which ALK inhibitors may be of utility.^9^ Other rare targetable activating fusions were present within the *CDKN2C*-null cohort, including in *BRAF, FGFR1*, and *NTRK1*.

If clinically indicated, this subtype could potentially be screened with genomic testing for *CDKN2C* loss, including through use of immunohistochemical surrogates.^56,57^ Well-established 1p/19q FISH testing may also play a role. Limitations of this study include its retrospective nature and the distinct population of patients highly enriched for aggressive tumors, mostly metastatic to distant sites. Given the highly selected, aggressive tumors examined in this study, the spectrum of smooth muscle neoplasms with *CDKN2C* loss may include lesions of benign and intermediate malignant potential not captured here. Additional studies will be needed to correlate the finding of *CDKN2C* homozygous loss in LMS with prognostic data and treatment outcomes. Comprehensive genomic profiling of LMS may provide insights into LMS biology and potentially inform therapeutic options, including specific cyclin-dependent kinase inhibitors.

## Previous presentation

None.

